# Urban ecology of arboviral mosquito vectors along the Kenyan coast

**DOI:** 10.1101/593350

**Authors:** Jonathan Karisa, Simon Muriu, Donwilliams Omuoyo, Boniface Karia, Doris Nyamwaya, Martin Rono, George Warimwe, Joseph Mwangangi, Charles Mbogo

## Abstract

**Background:** The emergence and re-emergence of arboviral infections particularly Chikungunya, dengue hemorrhagic fever, rift valley fever, and yellow fever in humans around the world threatens global health and security. The purpose of this study was to determine the urban ecology of the common arboviral mosquito vectors in urban Coastal Kenya areas.

**Materials and Methods:** The current study was conducted in urban settings of Kilifi and Mombasa counties in coastal Kenya in 2016 to 2017. Adult mosquitoes were collected both indoors and outdoors by CDC light traps and BG-Sentinel traps respectively. All blood fed mosquitoes were tested for blood meal sources by an Enzyme Linked Immunosorbent Assay (ELISA). Mosquito larvae were collected using standard dippers and pipettes. Egg survivorship in dry soil was evaluated by collecting of soil samples from dry potential breeding habitats, watering them for hatching and rearing of the eventual larvae to adults. Mosquitoes were screened for *Flavivirus, Alphavirus,* and *Phlebovirus* arboviruses using Reverse Transcriptase quantitative Polymerase Chain Reaction (RT qPCR).

**Results:** A total of 3,264 adult mosquitoes belonging to ten species of *Culex, Aedes* and *Anopheles* were collected. Overall, the predominant species were *Cx. quinquefasciatus* 72.4% (n=2,364) and *Ae. aegypti,* 25.7%, (n=838). A total of 415 breeding habitat types were identified indoors (n=317) and outdoors (n=98). The most productive habitat types in both indoors and outdoors were: assorted small containers, water tanks, drainages, drums and jericans. Overall, 62% (n=18) of the soil samples collected from the two sites (Kilifi and Malindi) were positive for larvae which were used as proxy to measure the presence of eggs. The mosquitoes had high preference for human blood (29.81%) and chicken (3.73%) but none had fed on either goat or bovine. Of 259 mosquitoes tested for viral infection, 11.6% were positive for *flavivirus* only.

**Conclusion:** Domestic and peri-domestic containers were identified to be the key breeding areas of arboviral vectors. Therefore, efforts should be put in place targeting the productive habitat types.

## Introduction

Arboviruses are arthropod borne viruses transmitted by an enormous number of haematophagous arthropod species, including but not limited to ticks, mosquitoes and sand flies (1). Mosquito borne viruses are responsible for serious viral disease outbreaks threatening human health and livelihoods especially dengue fever (2, 3), yellow fever (4), west Nile fever (5) and Chikungunya (6, 7). The emergence and re-emergence of arboviruses have significantly impacted on human and animal health as it results in global insecurity to all populations (35). They have been attributed to high level of morbidity and mortality particularly in sub-Saharan Africa and other tropical and subtropical environments (8). An estimated of 831 million people are living in an area at risk of at least one of arbovirus infections in the tropics and sub tropics region of the world (9). Hence, there exists a gap in knowledge and need of studies to increase the information and our understanding of the emergence and reemergence and danger to global health.

Rift valley fever virus (RVFV) which belongs to the family *Bunyaviridae,* genus *Phlebovirus* first was reported in 1912 (10). This vector borne virus is endemic in the tropics and sub tropics regions of Asia (11) and Africa (8). Several subsequent outbreaks of RVFV have been reported in different regions in Kenya (11, 13, 39, 40, 41, 42, 43).These outbreaks have resulted in loss of human and livestock lives in Kenya (16, 17). Dengue (family *Flaviviridae,* genus *flavivirus)* is endemic throughout much of Africa. This disease is often overshadowed by other febrile illness like malaria due to lack of awareness by health care providers, lack of diagnostic testing and surveillance (entomological and serological surveillances) (19). All four dengue viral strains/types have been reported, with outbreaks/epidemics being reported in almost the whole of African continent including the coastal Kenya, (17, 36). Dengue fever has been sporadic and has been reported in the north Eastern and Coastal Kenya where this is endemic (8, 16, 17, 44, 45, 46, 47). Chikungunya (family Togaviridae, genus *Alphavirus)* is another arboviral disease characterized by chills and arthralgia. Outbreaks Chikungunya fever have also been reported in west Africa (26, 27) and other parts of the continent including East Africa and Coastal Kenya (7,16,48,49,50). Other crucial arboviral infections which have been reported in Kenya includes: Yellow fever (51,52) and West Nile (16, 47,53,54). Kwale, Mombasa and Tana River Counties have reported significantly high levels of IgG antibodies against YFV and WNV (16,54). Studies have demonstrated the significant role of mosquitoes in maintenance of these arboviruses in nature during dry season through vertical transmission (41,57). Despite the long history of these infections, the epidemiology and public health effect in the coastal region of Kenya is still poorly understood. Therefore, detection of the arboviruses in the local vector, animal and human population hosts through active surveillance would constitute crucial components for effective control of unforeseen outbreaks.

Although there are over 300 species of mosquitoes that have been incriminated in arboviral transmission (1), *Aedes* and *Culex* mosquitoes have been blamed for transmission of 115 and 100 types of viruses respectively (1). Key species in arboviral transmissions are *Ae. Aegypti* (3) and *Culex quinquefasciatus* (36). *Aedes aegypti* has been responsible for the outbreak of several arboviruses in America (35, 36), West Africa (39) and Asia (40). On the other hand, *Cx. quinquefasciatus* have been responsible for the transmission of arboviruses in America (41), Asia (42), West Africa (43) and some parts of Europe (44). Previous studies in Kenya (12,13) have shown a high abundance of these arboviral vectors and a wide distribution along the Kenyan Coast with a clear temporal variation (8,15,18). However, there is limited routine entomological surveillance and current understanding of the ecology of the arboviral mosquito-vectors in urban coastal landscape.

There exists a high diversity and widespread distribution of arboviral mosquito vectors large due to occurrence of ideal breeding habitats (3). *Aedes aegypti* readily breeds in stagnant water especially in containers such as discarded plastic containers, bottles, coconut husks, old tires, drums, barrels, water storage tanks, obstructed roof gutters and broken bottles fixed on walls in and around human habitations(8, 15, 19). The occurrence of these breeding habitats in and around human habitations is indicative of the species adaption to domestic settings mostly living in close proximity to humans, preferentially and frequently feeding on them (49). Recent studies reported *Ae*. *aegypti* to breed mostly outdoors, in what may be a novel adaptive strategy by this vector, which traditionally is considered to have adapted to the domestic environment and breed mainly in indoor water storage containers (50). Human activities (water storage, use and disposal of water-holding containers and unplanned urbanization greatly influence *Ae*. *aegypti* breeding in individual households in urban settings (51). There are several key factors that significantly influence the productivity of *Ae*. *aegypti* in different container types including; the frequency of water replenishment, the availability of food for the larva, the degree of sunlight exposure and container covering (50–52). Due to unplanned urbanization and poor disposal of containers such as tyres, the outdoor environment provides more ideal environments as breeding areas due to the availability of numerous rain-filled discarded containers (51). Furthermore, *Aedes* mosquitoes eggs has ability to survive/overwinter/aestivate through dry periods a few centimetres down in moist soil/dry soils for several years (21,22) or even if the container is refilled. Detection of eggs in dry soils from potential breeding sites (tyres, water tanks, tree holes, assorted small containers etc.) provides more reliable information the breeding preferences selected by gravid females (56) and other ecological adaptive strategies of these vector species. Hence, the development and implementation of vector intervention requires clear understanding of *Aedes aegypti* ecology and plasticity in its breeding habitats and behaviour. The inclusion and derivation of populations estimates from eggs samples from both natural and artificial potential breeding habitats is crucial in evaluating the larval ecology and population dynamics of the species.

*Culex* mosquitoes are the most common species of mosquitoes with liberal/diverse breeding habitats. Although there are more than 700 species of Culex mosquitoes with diverse behavioral and adaptive characteristics, *Cx. quinquefasciatus* is the most dominant and widespread species that has also been incriminated in several pathogens transmission. The species is involved in transmission of West Nile virus, Japanese encephalitis, St. Louis encephalitis, chikungunya, Rift valley fever virus, filariasis and avian malaria (14, 25, 26, 27, 28). Their involvement in transmission of these diverse groups of pathogens is due to its mixed/liberal blood feeding behavior (blood meal sources) that range from rodents, reptiles avian, primates and humans (62). *Culex quinquefasciatus* mosquitoes have been shown to breed/oviposit in water with high organic content mostly in rice paddies, canals, swimming pools, chambers, drainage, rain pools, ditches, rock pools, septic tanks, tree holes and run off from agricultural treatment plants (29,30,31,32). Due to the rapid growth and development of urban areas in tropics and the involvement of *Cx. quinquefasciatus* in the transmission arboviruses, this mosquito has become a matter of growing concern in recent years (67). There is need for the development of an effective vector control programme or strategy against this species. This will ultimately require knowledge of some aspects of its ecology. Thus, comprehensive information on the bionomics of the target species is essential before implementation of any control program. Therefore, determining the ecology of these arboviral vectors would provide a way forward in terms of pesticide application and control of arboviral infections.

Intervention strategies against arboviruses is mainly based on reduction of the vector species population and subsequent reduction in vector-human contact (58,59). Development of effective vaccines against arboviral infection like Yellow fever in humans (70) and rift valley fever in livestock (71) has been crucial in preventing arboviral infections. However, vaccination programs are constrained by the limited number of effective vaccines for a majority of circulating arboviruses (72). Consequently, the most effective alternative is to focus on practical procedures to monitor the vector populations and their interaction with human host thereby reducing risks of human exposures to arboviruses. Mosquito vectors control is done through combination of several strategies in a synergistic manner through integrated vector management programs involving pesticide application, public education and biological control in order to reduce the potential for disease transmission and biting nuisance (57,58). The current study was undertaken with overarching aim of elucidating information/knowledge of the ecological parameters/aspects of arboviral vectors in urban tropical coastal settings. The goal was to generate information essential arboviral transmission risks and development of intervention strategies against their mosquito vector populations.

## Materials and Methods

### Study area

The study was conducted in three urban coastal areas of Mombasa (4.0435°S; 39.6682°E), Kilifi (3.5107° S; 39.9093° E) and Malindi (3.2192° S; 40.1169° E) in Kenya from November 2016 to April 2017. The Kenyan Coastal region is characterized by dense forests, savanna type of vegetation, seasonal swamps, dry thorn bushes and diverse plantations interspersed with furrow land. Sisal, coconut, and cashew nut plantations are prominent along the coast although subsistence farming is common in inland areas. Altitudes range from 0-400 meters above sea level. The region experiences a bimodal form of rainfall with the long monsoon rains occurring in April to July and the short rains occurring between October and December. The relative humidity ranges from 55% to 65% and temperature from 20°C to 35°C with an annual rainfall of 750 to 1,200 mm. The two counties of Mombasa and Kilifi are characterized by flat topography. The rural areas mainly inhabited by the Mijikenda and Swahili communities while urban areas have mixed population of different Kenyan communities and tourists from around the globe. The major economic activities are: tourism, fishing, commercial trade and retail, and service professions whereas the informal economic sector is comprised of street vendors, sex workers, and tour guide services. These are interspersed with commercial, undeveloped, farmed, and residential areas. Houses are mainly made of concrete or mud walls and iron sheet or palm leaf (Makuti), roofing Most households keep goats, chickens, cats, ducks, dogs and cattle. In each site, three residential estates were selected for larval and adult mosquito sampling.

### Mosquito larval sampling and habitat characterization Larval sampling

Mosquito larval sampling was done in three randomly selected residential estates in each of the three urban study sites of Mombasa, Kilifi and Malindi. Residential estates are housing estates designed as autonomous suburb with a centred small commercial center. The urban estates sampled for larvae in Mombasa county included Port reitz, Bombolulu and Tudor while in Kilifi, Bofa, Mnarani and Mtaani were selected and in Malindi, it included Kisumu ndogo, Ngala and Complex area. During the study, all potential *Aedes* and *Culex* mosquito breeding habitats inside houses and peridomestic locations were identified and checked mosquito larvae. The indoor and outdoor water containers were identified and visually checked for mosquito larvae and pupae. Depending on the habitat size and type, mosquito sampling was done using standard dippers (350ml) 5-20 dippers per containers depending on size or census was done by pipetting. The mosquito samples from each habitat were placed in individually labeled whirl paks, placed in a cooler box and transferred to the laboratory for further processing.

### Larval habitat classification

Drums were defined as cylindrical containers of capacity between 50-200-liter while water tanks were defined as any water storage container with 200-1000 liters of water storage capacity. The assorted small containers comprised small plastic/metallic containers of less than 10liters water holding capacity.

### Sampling for *Aedes aegypti* egg survivorship in dry substrate

Sampling for *Aedes* eggs was conducted in all potential *Aedes* breeding habitats identified in the residential estates in Kilifi and Malindi urban study sites. During sampling, the dry soil or substrate from the identified potential breeding habitats were sampled by scooping a handful of the soil or substrate with a spatula and placed in while paks and transferred to the laboratory for further processing in the KEMRI insectary.

### Adult mosquito sampling

Adult mosquitoes were sampled using Biogent (BG) Sentinel trap and CDC light traps. The BG-Sentinel traps were primarily deployed for surveillance of adult *Aedes aegypti* mosquitoes as describe by Maciel-de-Freitas (75). A total of nine (9) BG sentinel traps baited with carbon dioxide (dry ice) were randomly set outdoors from 6:00 am to 5:00 pm, in each of the three residential estates in three-urban setting. The traps were set systematically on the ground at intervals of 100m from each other and sampling was done only once in each of the representative residential estates.

The CDC light traps were set up between 1700-1830 hours and left to run throughout the night and collected at 06.00hrs the following morning. Forty (40) randomly selected houses were sampled using light traps in Mombasa (Bombolulu=10, Tudor=10 and port reitz=20). In Malindi the 40 traps included 10 traps in Kisumu ndogo, 10 traps in Complex area and 20 traps in Ngala) while the 30 traps in Kilifi were equally in Bofa, Mtaani and Mnarani. The mosquito samples collected were transferred to the laboratory for further processing in a cool box.

### Laboratory sample (mosquito) processing

#### Egg processing

In the insectary, the habitats soil or substrate sample collections were placed in individually labeled basins and one litre (1L) of de-ionized water added and allowed to settle. The basins were monitored daily for eggs-to hatch into first instar larvae. The resultant larvae were reared to pupae that were enumerated, recorded and transferred to pupal cages for adult emergence. All the soil/substrate samples were monitored for two (2) weeks and those that did not register any larvae were regarded as negative samples after this period. The emergent adults were enumerated and morphotyped using identification keys.

### Larval rearing

In the laboratory, immature mosquito samples collected from the different habitats in the field were sorted and categorized into Anopheline, Culicine or Aedine larvae. The larvae were further grouped as early (L1 and L2) or late (L3 and L4) and reared in labeled plastic basin using water obtained from the site of larval collection. Larval development was monitored daily and all pupae harvested using a pipette, placed in pupal cups in mosquito cages for adult emergence. Emerging adult mosquitoes were maintained alive with 10% glucose concentration until identification. Temperature for larvae breeding and rearing was maintained at between 32-34°C and in the adult breeding room at between 26-28°C while relative humidity of 70-80% for both larvae and adults.

### Adult mosquito processing and identification

All adult mosquitoes from the field were killed by freezing the BG sentinel and light trap contents at −20°C for 2 minutes. The samples were then sorted from other arthropods, sexed and morphologically identified to species level using identification keys by Edwards (62). All the samples were preserved in 1.5 ml cryogenic vials in −80°C for arbovirus testing

### Blood meal analysis

All blood fed mosquitoes from the field collected adult mosquitoes were carefully cut transversely at mid-section to separate head and thorax section from the abdomen. The abdominal section was placed in a labeled vial while the rest were preserved appropriately for arboviral testing. Blood meal analysis was done using Enzyme Linked Immunosorbent Assay (ELISA) method as described previously (60, 61). Results were read visually through colour change (homogenous greenish blue colour for positive and clear for negative samples).

### RNA extraction

All adult mosquito samples from larval, habitat substrate or soil, and adult collections were processed on chill table by preparing mosquito (1 to 25 mosquitoes per pool) by site, method of collection, species and sex in preparation for arboviral analysis. If not processed immediately all the samples were preserved in 1.5 ml cryogenic vials in −80°C. RNA was extracted from mosquito samples using Trizol^®^-LS – Chloroform extraction method (78). Briefly, 1 ml of TRIzol™ reagent was added to the sample (pools of whole mosquitoes) and then homogenized followed by addition of 0.2 ml chloroform to the homogenate and vortexed for 30 seconds. The resultant homogenate was incubated for 2-3 minutes then centrifuged at 12,000rpm for 15 minutes at 4 °C. The aqueous phase was then transferred to a fresh eppendorf tube and the RNA precipitated by mixing with 0.5 ml isopropanol followed by incubation at room temperature for 10 minutes. The mixture was then centrifuged at 12,000rpm for 10 minutes at 4°C and the supernatant removed before washing the pellets with 1 ml of 75% ethanol by flicking followed by centrifugation at 7,500rpm for 10 minutes at 4°C. The supernatant is removed and the pellets air-dried. The final RNA pellet was dissolved in 50 μl of nuclease-free water at room temperature and stored on ice or frozen at −80°C ready for screening.

### Arbovirus screening

The extracted RNA was tested using primers targeting *Flavivirus, Alphavirus,* and *Phlebovirus* arboviral genera. Virus detection and amplification was done using the QuantiFast Multiplex RT-PCR + R kit (Qiagen) in conjunction with primers and probes designed for generic amplification of Flavivirus non-structural 5 gene (NS5), alphavirus non-structural protein 4 (NSP4) gene and Phlebovirus primers targeting the Large (L) and small (S) segments. The protocol for Flavivirus, alphavirus and Phlebovirus assay have been described elsewhere (70, 71, 72). Dengue specific assay was performed to all samples that tested positive for Flavivirus. The ABI 7500 real time PCR (Applied Biosystems, USA) was used for amplification.

### Data management and analysis

Data collected was entered into Microsoft excel and analyzed in Stata statistical package (82).The mean number of immature mosquitoes in indoors and outdoors was calculated and the difference compared within each site using a t-test. Chi square was used to measure the association between site, sex and species variation with regards to flavivirus positivity. Statistical differences between and among groups was deemed significant at p < 0.05.

The larval mosquito infestation indices were calculated as House Index (HI)—the percentage of houses positive with immature mosquitoes, Container Index (CI)—the percentage of water holding containers in which mosquito breeding is occurring and Breteau Index (BI)—the number of positive containers per 100 houses. The following formulae were used to determine these indices:

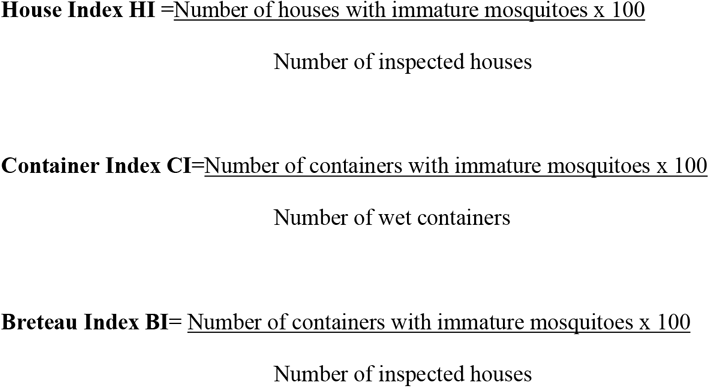

Shannon diversity index (*H*) was used to characterize species diversity in the three study sites in the urban coastal Kenya. Shannon’s index takes into consideration for both abundance and evenness of the species present. The proportion of species (*i*) relative to the total number of species (*pi*) was calculated and then multiplied by the natural logarithm of this proportion (ln*pi*). The resulting product, which is always a negative value, was summed across species and multiplied by −1

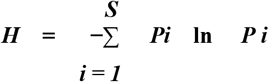

Shannon’s equitability (*EH*) was calculated by dividing *H* by Hmax (where Hmax = ln*S*, the total number of species in the community (richness). Equitability/evenness deduce a value between 0 and 1 with 1 being complete evenness.

## Results

### Larval habitats diversity and productivity

A total of 415 mosquito breeding habitats were identified both inside (317) and outside (98) houses. Out of these, 168 (40.5%) were found in Kilifi, 114 (27.5%) in Malindi and 133 (32.04%) in Mombasa (Table 1). Fourteen different larval habitats were identified and sampled in the three study sites. Overall, the most prevalent breeding habitat in the three sites were Jericans (66.9%), followed by water tanks (10.6%), small containers (6.75%) and drainage channels (6.02%). Other habitats are: buckets, basins, ditches, water troughs, flower pots, swimming pools, chambers and earthen water pots (Table 1). In Kilifi, seven different types of mosquito-breeding habitats were identified. The most abundant habitat types were jericans (61.90%), followed by water tank (16.07%), assorted small containers (11.31%), drainages (6.55%), and tyres (1.79%). Others that were reported in small numbers include ditches and drainage chambers. Similarly, in Malindi, eleven different habitat types were identified (Table 2). The most abundant habitat type was jericans (71.05%) followed by small containers (7.02%), water tanks (5.26%), drainages (5.26%) and tyres (0.88%). Other habitats encountered though in small numbers include bucket, basin, ditch water trough, flowerpot and swimming pool (Table 2). In Mombasa, ten different habitats were surveyed and identified; jericans (69.92%) were the most abundant habitat type followed by water tanks (8.27%), drainages (6.02%), drums (5.26), and tyres (3.76%). Other habitat types that were encountered though in smaller quantities include: assorted small containers, bucket, basin, ditch and water pot (Table 1).

**Table 1.** Summary of the habitat productivity for indoor and outdoor locations in the three sites of urban coastal Kenya

| Kilifi |  |  |  |  |  |  | Malindi |  |  |  |  |  | Mombasa |  |  |  |  |  |
| --- | --- | --- | --- | --- | --- | --- | --- | --- | --- | --- | --- | --- | --- | --- | --- | --- | --- | --- |
| Habitat Type | Indoor |  |  | Outdoor |  |  | Indoor |  |  | Outdoor |  |  | Indoor |  |  | Outdoor |  |  |
|  | No of Habitats (+ ve habitats) | Mean Larvae | Mean Pupae | No of Habitats (+ ve habitats) | Mean Larvae | Mean Pupae | No of Habitats (+ ve habitats) | Mean Larvae | Mean Pupae | No of Habitats (+ ve habitats) | Mean Larvae | Mean Pupae | No of Habitats (+ ve habitats) | Mean Larvae | Mean Pupae | No of Habitats (+ ve habitats) | Mean Larvae | Mean Pupae |
| Small container | 9 (1) | 10 | 0 | 10 (5) | 108 | 46 | 0 (0) | 0 | 0 | 8 (2) | 27 | 0 | 0 (0) | 0 | 0 | 1 (0) | 0 | 0 |
| Drum | 0 (0) | 0 | 0 | 0 (0) | 0 | 0 | 0 (0) | 0 | 0 | 0 (0) | 0 | 0 | 7 (6) | 43 | 3 | 0 (0) | 0 | 0 |
| Water tank | 24 (6) | 9 | 1 | 3 (1) | 15 | 0 | 3 (0) | 0 | 0 | 3 (2) | 74 | 70 | 9 (3) | 44 | 5 | 2 (1) | 41 | 0 |
| Jerican | 104 (2) | 7 | 1 | 0 (0) | 0 | 0 | 62 (1) | 26 | 4 | 19 (1) | 50 | 0 | 91 (6) | 28 | 1 | 2 (0) | 0 | 0 |
| Bucket | 0 (0) | 0 | 0 | 0 (0) | 0 | 0 | 0 (0) | 0 | 0 | 3 (2) | 58 | 50 | 2 (0) | 0 | 0 | 0 (0) | 0 | 0 |
| Basin | 0 (0) | 0 | 0 | 0 (0) | 0 | 0 | 0 (0) | 0 | 0 | 1 (0) | 0 | 0 | 3 (0) | 0 | 0 | 0 (0) | 0 | 0 |
| Drainage channels | 2 (0) | 0 | 0 | 9 (3) | 87 | 4 | 0 (0) | 0 | 0 | 6 (0) | 0 | 0 | 0 (0) | 0 | 0 | 8 (1) | 17 | 4 |
| Ditch | 0 (0) | 0 | 0 | 1 (0) | 0 | 0 | 0 (0) | 0 | 0 | 1 (0) | 0 | 0 | 0 (0) | 0 | 0 | 2 (0) | 0 | 0 |
| Tyre | 0 (0) | 0 | 0 | 3 (0) | 0 | 0 | 0 (0) | 0 | 0 | 1 (1) | 20 | 0 | 0 (0) | 0 | 0 | 5 (1) | 30 | 0 |
| Water trough | 0 (0) | 0 | 0 | 0 (0) | 0 | 0 | 0 (0) | 0 | 0 | 3 (3) | 36 | 0 | 0 (0) | 0 | 0 | 0 (0) | 0 | 0 |
| Flower pot | 0 (0) | 0 | 0 | 0 (0) | 0 | 0 | 0 (0) | 0 | 0 | 3 (0) | 0 | 0 | 0 (0) | 0 | 0 | 0 (0) | 0 | 0 |
| Swimming pool | 0 (0) | 0 | 0 | 0 (0) | 0 | 0 | 0 (0) | 0 | 0 | 1 (0) | 0 | 0 | 0 (0) | 0 | 0 | 0 (0) | 0 | 0 |
| Chambers | 0 (0) | 0 | 0 | 3 (0) | 0 | 0 | 0 (0) | 0 | 0 | 0 (0) | 0 | 0 | 0 (0) | 0 | 0 | 0 (0) | 0 | 0 |
| Water pot | 0 (0) | 0 | 0 | 0 (0) | 0 | 0 | 0 (0) | 0 | 0 | 0 (0) | 0 | 0 | 1 (0) | 0 | 0 | 0 (0) | 0 | 0 |
| **Total** | 139 (9) | 26 | 2 | 29 (9) | 210 | 50 | 65 (1) | 26 | 4 | 49 (10) | 216 | 120 | 113 (15) | 115 | 9 | 20 (3) | 88 | 4 |

**Table 2.** Indoor site-specific House, Container and Breteau indices for the 3 coastal urban area

| Sampled site | No. houses of sampled houses | No. of positive houses | HI | No. of wet habitats | No. of positive habitats | CI | BI |
| --- | --- | --- | --- | --- | --- | --- | --- |
| **Kilifi** | 30 | 7 | 23.33 | 139 | 9 | 6.47 | 30.00 |
| **Malindi** | 11 | 1 | 9.09 | 65 | 1 | 1.54 | 9.09 |
| **Mombasa** | 14 | 10 | 71.43 | 113 | 15 | 13.27 | 107.14 |
| **Overall** | **55** | **18** | **32.73** | **317** | **25** | **7.89** | **45.45** |

Overall, the most productive containers (habitats) indoors were drums, small containers, jericans and water tanks whereas for outdoors the most productive containers were drainage channels, small containers, tyres, water tanks, jericans and water troughs (Figure 2). There was a significant association between habitat type and immature productivity (χ^2^ (df= 13, N=415) = 134.1488, p<0.001). A total of 18 breeding habitats in Kilifi, (6 *%* indoors, 31%outdoors) were positive for mosquito immature stages. There was significant difference in the density of immatures between indoor and outdoor (P< 0.05). The most productive indoor containers in Kilifi were small containers, water tanks and jericans whereas outdoors were drainage channels, small containers and water tanks (Figure 2). In Malindi, 12 habitats (2% indoors, 22% outdoors) were found to be positive for mosquito immatures and there was no significant difference between them (P>0.05). The most productive indoor habitats in Malindi were only jericans. On the other hand, the most productive containers outdoors were water tanks, jericans, others, small containers and the least were tyres (Figure 2). In Mombasa, 18 habitats (13% indoors, 15% outdoors) were found to be positive for mosquito immatures (Figure 2). Similarly, there was no significant difference in the density of immatures between indoor and outdoor (P>0.05). In Mombasa, the most productive indoor habitats were water tanks, drums and jericans whereas for outdoor habitats, water tank was the most productive habitat type, followed by tyres and the least were drainage channels (Table 1).

**Figure 1.**
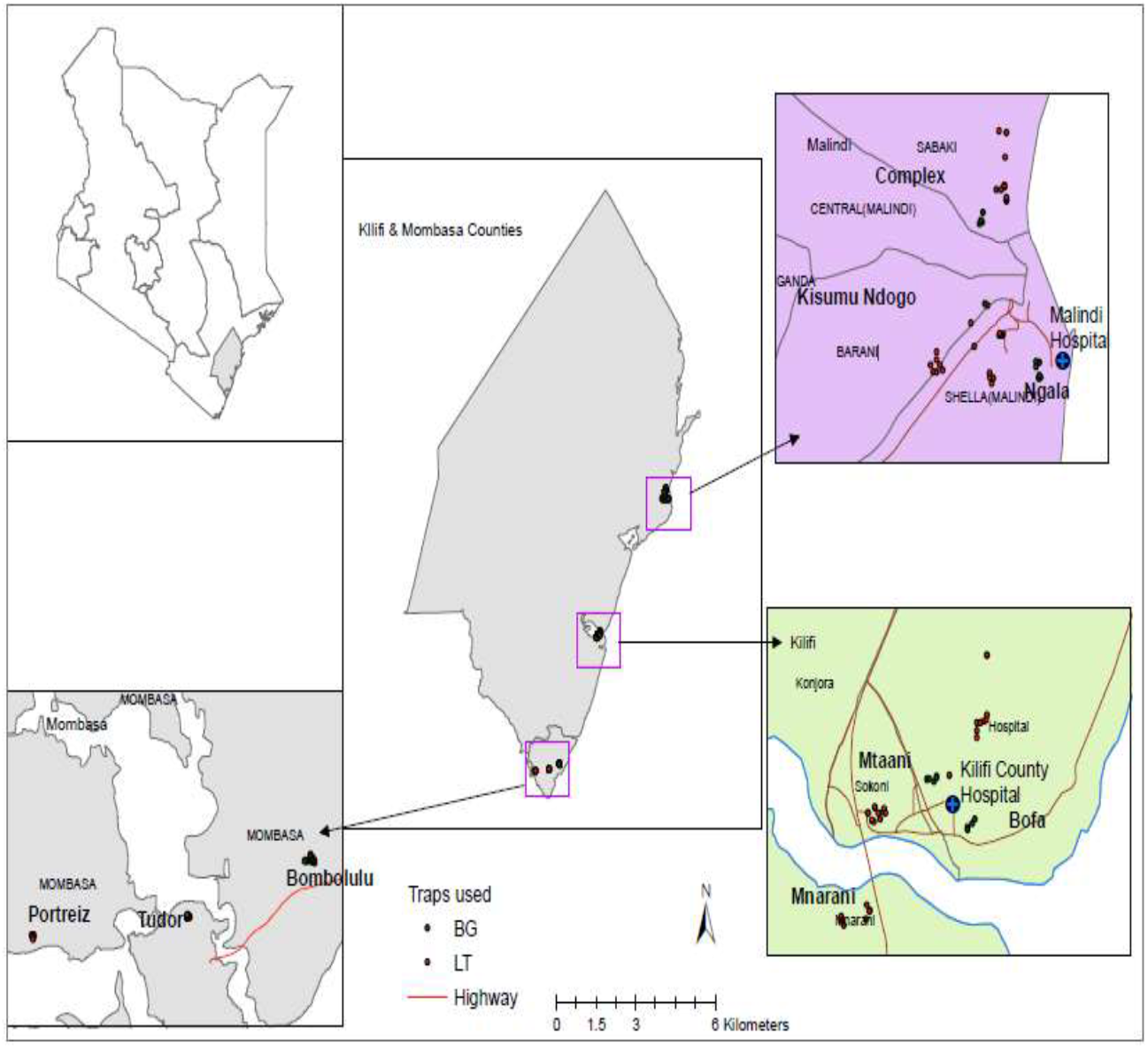
Map of the Mombasa and Kilifi Counties showing the position of sites from which samples were collected

**Figure 2.**
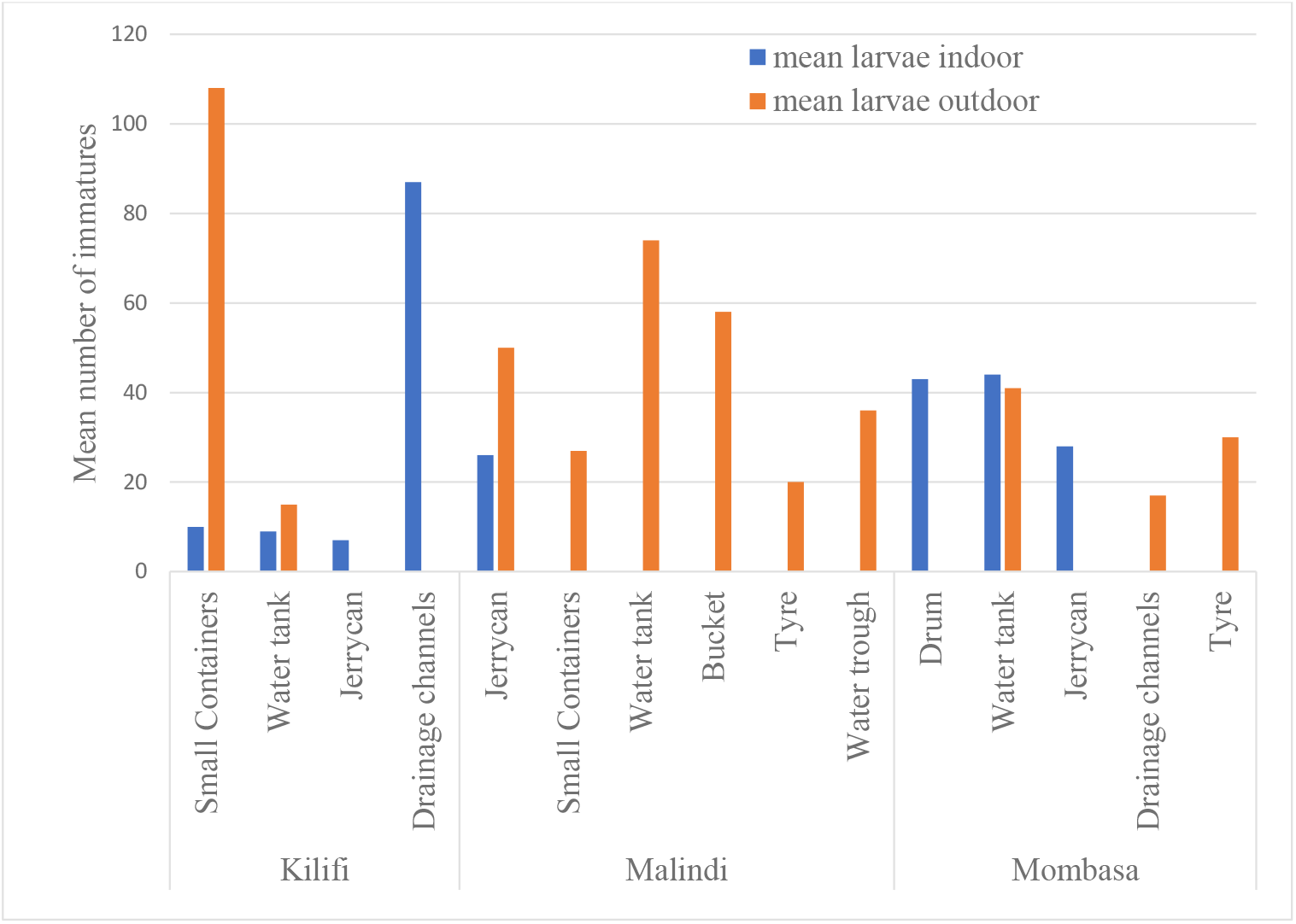
The most productive containers indoors and outdoors in Kilifi, Malindi and Mombasa sites

### Species composition in larval habitat collection

Overall, 889 adult mosquitoes four species belonging to two genera *(Aedes and Culex)* emerged from the larval population collected. Of the four species that emerged. Majority were *Aedes aegypti* (85.3%) wand the rest being *Culex quinquefasciatus* (12.60%), *Ae. vittatus* (1.12%), and *Cx. zombiensis* (1.01%). Indoor immatures resulted in purely and exclusively *Ae. aegypti* mosquitoes whereas as outdoor had both *Ae. aegypti* and *Cx. quinquefasciatus* mosquitoes.

### Larval infestation indices

Fifty-five (55) houses were sampled from the three sites for mosquito habitats. Out of these houses, 18 had containers that were positive for Ae. *aegypti* immatures, giving an overall HI of 32.72%. A total of 317 containers were inspected indoors giving an overall CI of 8% and Breteau index of 45.45. Mombasa had the highest indices (HI of 71.43, CI of 13.27 and BI of 107.14) compared to Malindi and Kilifi (Table 2).

### Mosquito egg survivorship in dry habitats

A total of 29 dry habitat substrate/ dry habitat soil samples were collected from water tank (n=2), small container (n=1), tyres (n=16) and flower pots (n=10). Overall, 62% (n=18) of the soil samples collected from the two sites (Kilifi and Malindi) were positive for larvae. Five hundred and six (506) adult mosquitoes resulted from the larvae reared from the dry breeding habitats substrate. three *Aedes* species were identified including *Aedes aegypti* (98.4%), *Aedes hirsutus* (1.4%) and *Aedes vittatus* (0.2%) (Table 3).

**Table 3.** Summary of the soil samples collected from different container, positive habitats and the mosquito species that emerged

| Site | Habitat type | No. of habitats ( % +ve) | Mosquito species | Total adults Emerged |
| --- | --- | --- | --- | --- |
| **Kilifi** | Tyre | 7 (57) | *Ae. aegypti* | 319 |
|  |  |  | *Ae. hirsutus* | 7 |
|  |  |  | *Ae. vittatus* | 1 |
|  | Flower pot | 3 (66) | *Ae. aegypti* | 2 |
|  | Water tank | 2 (0) | 0 | 0 |
| **Malindi** | Small container | 1 (100) | *Ae. aegypti* | 16 |
|  | Tyre | 9 (77) | *Ae. aegypti* | 79 |
|  | flower pot | 7 (71) | *Ae. aegypti* | 82 |

### Adult mosquito distribution and abundance collections

The relative abundance of adult mosquitoes collected indoors and outdoors by the Biogents Sentinel (BG) traps and Light traps (LT) is summarized in table 4. Overall, 3,264 mosquitoes belonging to three genera *(Culicines, Aedes* and *Anopheles)* and 10 species were collected. *Aedes Aegypti* (838) and *Cx. quinquefasciatus* (2,364) were the most common species, and the least were *Ae. mcintoshi, Ae. pembaensis,* and *Cx. Annulioris,* (n=1). *Cx. quinquefasciatus* were mostly collected indoors (n=2,140) compared to outdoors (n=260) while more *Ae. aegypti* mosquitoes were captured outdoors (n=816) compared to indoors (n=22) (Table 4). The Shannon diversity index (H) and evenness *(EH)* of mosquito species indicated a higher species diversity in Kilifi *(H* =0.840) compared to Malindi (H = 0.662) and Mombasa (H =0.385). Similarly, mosquitoes were evenly distributed in Kilifi *(EH* =0.469) compared to Malindi *(EH* =0.370) and Mombasa *(EH* =0.215).

**Table 4.**
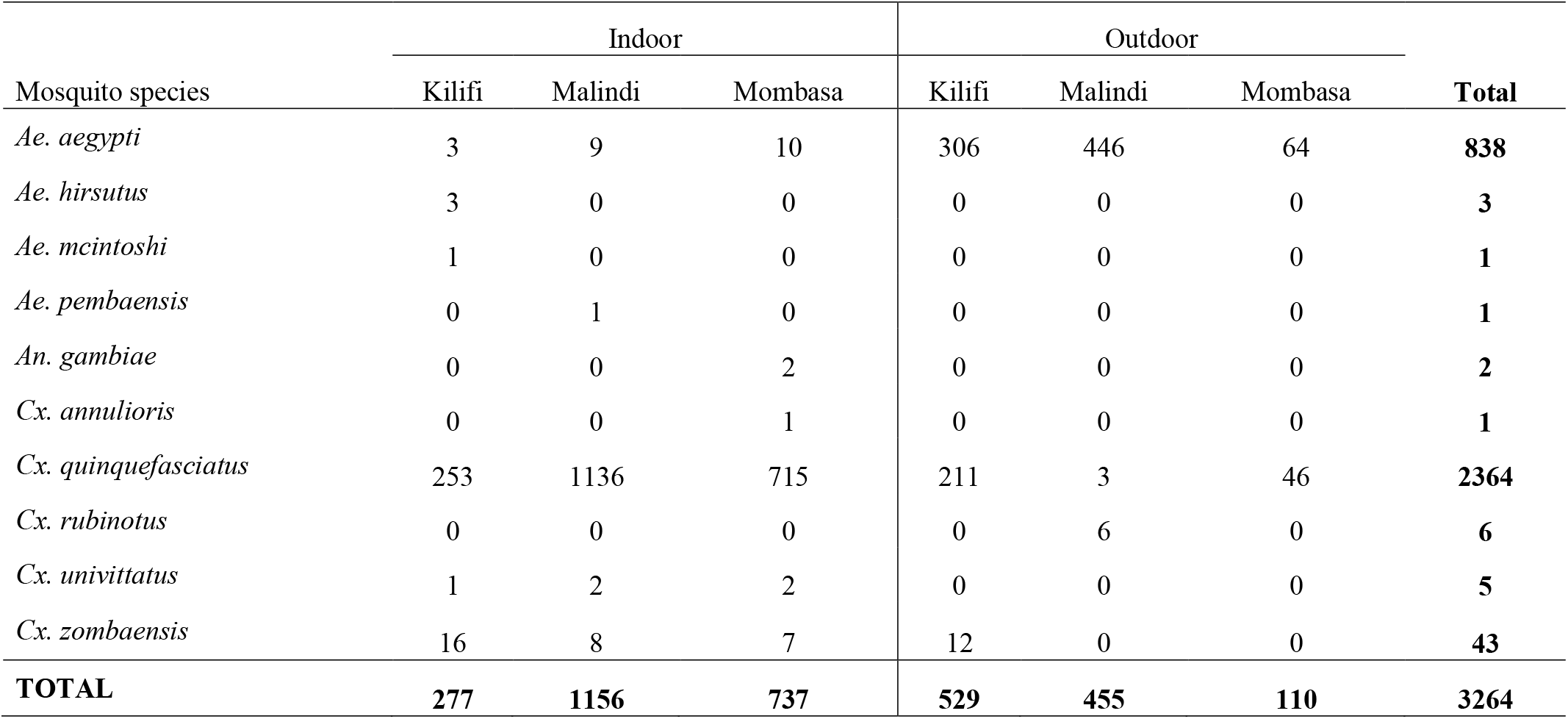
Relative abundance of mosquito species collected in the three urban areas of coastal Kenya

| Mosquito species | Indoor |  |  | Outdoor |  |  | Total |
| --- | --- | --- | --- | --- | --- | --- | --- |
|  | Kilifi | Malindi | Mombasa | Kilifi | Malindi | Mombasa |  |
| *Ae. aegypti* | 3 | 9 | 10 | 306 | 446 | 64 | **838** |
| *Ae. hirsutus* | 3 | 0 | 0 | 0 | 0 | 0 | **3** |
| *Ae. mcintoshi* | 1 | 0 | 0 | 0 | 0 | 0 | **1** |
| *Ae. pembaensis* | 0 | 1 | 0 | 0 | 0 | 0 | **1** |
| *An. gambiae* | 0 | 0 | 2 | 0 | 0 | 0 | **2** |
| *Cx. annulioris* | 0 | 0 | 1 | 0 | 0 | 0 | **1** |
| *Cx. quinquefasciatus* | 253 | 1136 | 715 | 211 | 3 | 46 | **2364** |
| *Cx. rubinotus* | 0 | 0 | 0 | 0 | 6 | 0 | **6** |
| *Cx. univittatus* | 1 | 2 | 2 | 0 | 0 | 0 | **5** |
| *Cx. zombaensis* | 16 | 8 | 7 | 12 | 0 | 0 | **43** |
| **TOTAL** | **277** | **1156** | **737** | **529** | **455** | **110** | **3264** |

### Blood meal sources

Out of the 161 blood fed female mosquitoes tested by ELISA for host blood meal sources, 91%, were from *Culex* (n= 146) and the rest *Aedes* (9%, n=15) species (Table 5). The samples were tested for against four blood mea source/antisera namely: bovine, chicken goat and human. Overall, majority of the blood meals identified were of human origin (29.81%) chicken (3.73%) and none from goat or bovine although 66.46% could not be identified (Table 5). The mosquitoes analyzed comprised of *Culex quinquefasciatus* (n=143), *Aedes aegypti* (n=15) and *Cx. univittatus* (n=3) (Table 5). Overall, the blood meal preference varied between outdoor and indoor samples with majority at indoor locations (n= 140) and to a lesser extent on outdoors (n=21). Larger percentage of the indoor mosquitoes had fed on human (n = 44), chicken (n = 4) and the rest had fed on unspecified hosts (n = 92), whereas for the outdoor collected samples, majority had fed on humans (n = 6), chicken (n = 2) and the rest had fed on unspecified hosts (n =13).

**Table 5.** Blood meal sources of the blood fed mosquitoes collected in Malindi, Kilifi and Mombasa urban areas

| Species | Site | Location | No. tested | Human (%) | Bovine (%) | Goat (%) | Chicken (%) | Unidentified (%) |
| --- | --- | --- | --- | --- | --- | --- | --- | --- |
| *Ae. aegypti* | Kilifi | outdoor | 4 | 3 (75) | 0 (0) | 0 (0) | 0 (0) | 1 (25) |
|  |  | indoor | 1 | 0 (0) | 0 (0) | 0 (0) | 0 (0) | 1 (100) |
|  | Malindi | outdoor | 9 | 0 (0) | 0 (0) | 0 (0) | 0 (0) | 9 (100) |
|  | Mombasa | indoor | 1 | 1 (100) | 0 (0) | 0 (0) | 0 (0) | 0 (0) |
| *Cx. quinquefasciatus* | Kilifi | outdoor | 7 | 2 (28.6) | 0 (0) | 0 (0) | 2 (28.6) | 3 (42.9) |
|  |  | indoor | 19 | 11(57.9) | 0 (0) | 0 (0) | 1 (5.3) | 7 (36.8) |
|  | Malindi | indoor | 73 | 22(30.1) | 0 (0) | 0 (0) | 3 (4.1) | 48 (65.8) |
|  | Mombasa | outdoor | 1 | 1 (100) | 0 (0) | 0 (0) | 0 (0) | 0 (0) |
|  |  | indoor | 43 | 7 (16.3) | 0 (0) | 0 (0) | 0 (0) | 36 (83.7) |
| *Cx. univittatus* | Malindi | indoor | 1 | 0 (0) | 0 (0) | 0 (0) | 0 (0) | 1 (100) |
|  | Mombasa | indoor | 2 | 1 (50) | 0 (0) | 0 (0) | 0 (0) | 1 (50) |
| **Total** |  |  | **161** | **48(29.8)** | **0 (0)** | **0 (0)** | **6 (3.7)** | **107(66.5)** |

Majority of the *Culex quinquefasciatus* mosquitoes which fed on human (n= 40) and chicken (n=4) were captured indoor and to lesser extent in the outdoor environs humans (n=3) and chicken (n=2) (Table 5). In Kilifi, 26 *Culex quinquefasciatus* were tested for host blood meal. Out of the 26 *Culex quinquefasciatus,* 50% (n=13) tested positive for human blood meal while 11.54% (n=3) had fed on chickens with 38.46% (n=10) unknown. *Culex quinquefasciatus* mosquitoes could have fed on humans in both indoors and outdoors with 57.9% and 28.6% respectively. This species could have also fed on chicken in both indoors and outdoors represented by 28.6% and 5.3% respectively. In Malindi, 73 *Culex quinquefasciatus* were tested for the host blood meal source. Out of the 73 *Culex quinquefasciatus,* majority had predominantly fed on humans (30.14%) indoors, while 4.11% had fed on chicken blood meals indoors and the rest (65.75%) had fed on unidentified blood meal sources. *Culex quinquefasciatus* fed on humans and chicken indoors. In Mombasa, 44 *Culex quinquefasciatus* were tested for blood meal host sources. Out of the 44 *Culex quinquefasciatus* mosquitoes, 18.18% (n=8) tested positive for human blood meal whereas unknown hosts sources constituted 84.09 (n=36) (Table 5).

For *Aedes aegypti,* 15 mosquitoes were tested for host blood meals. Eighty-seven percent 87% (n=13) were captured outside houses while 13% were trapped inside houses. Out of the 15 *Aedes aegypti,* 73% (n=11) had fed on unidentified hosts while the rest (27%, n=4) had feed on humans (3 outdoors & 1 indoors). In Kilifi, five *Aedes aegypti* were tested for host blood meal majority of which had predominantly fed on humans (60%, n=3) and the rest (40%, n=2) were fed on an unidentified host. *Aedes aegypti* was found to feed both indoors (20%, n=1) and outdoors (80%, n=4). For the outdoor (n=3), preferentially fed on human while one fed on unidentified hosts. For the case of indoors, these mosquitoes fed on unidentified or unknown hosts. In Malindi, nine *Aedes aegypti* were tested for the host blood meal source. None of them had fed on human, bovine, goat or chicken. *Aedes aegypti* mosquitoes exclusively fed outdoors on unidentified hosts. In Mombasa, one *Aedes aegypti* mosquito was tested for blood meal host sources and had fed on human blood indoors. A single (100%) *Ae. aegypti* mosquito tested from this area fed on human inside houses (Table 5).

Three *Cx. univittatus* were tested for host blood meal. All of these had sourced their blood meal inside houses. Majority (67%, n=2) of these had fed on unidentified hosts while the rest (33%, n=1) fed on humans. In Malindi, one *Cx. univittatus* was tested for the host blood meal source and had fed on unknown hosts inside house. In Mombasa, two *Cx. univittatus* were tested for blood meal host sources. Fifty percent (50%, n=1) of these species had fed indoors on human blood while rest 50% (n=1) had fed an unidentified host (Table 5).

### Arboviruses diversity in mosquitoes

Out of the 259 pools screened against the three viral genera and 11.58% pools tested positive (Table 6). Overall, the pools consisted of 129 *Ae. aegypti* pools and 130 *Cx. quinquefasciatus* pools. The overall positive pools (n=30) were only positive to *Flavivirus* and none for either *Phlebovirus* or *Alphavirus.* The *Aedes aegypti* had a significantly higher *(^χ2^, (df=1, n=30)* = 18.4398, P=0.000) proportion of virus positive pools (87%, n=26) compared to *Cx. quinquefasciatus* (13%, n=4).

**Table 6.** Total number of pools positive for flavivirus in the Kilifi, Mombasa and Malindi

| Species | Site | Sex | Total number of pools (positive) |  |  |  |
| --- | --- | --- | --- | --- | --- | --- |
|  |  |  | BG | LT | Larvae | soil sample |
| *Ae. aegypti* | Kilifi | F | 8 (3) | 1 (1) | 5 (1) | 5 (0) |
|  |  | M | 14 (5) | 4 (3) | 6 (2) | 8 (3) |
|  | Malindi | F | 13 (3) | 3 (0) | 6 (0) | 6 (2) |
|  |  | M | 11 (0) | 2 (0) | 7 (0) | 4 (0) |
|  | Mombasa | F | 4 (1) | 3 (0) | 5 (0) | 0 (0) |
|  |  | M | 3 (0) | 3 (1) | 8 (1) | 0 (0) |
| Sub total |  |  | 53 (12) | 16 (5) | 37 (4) | 23 (5) |
| *Cx. quinquefasciatus* | Kilifi | F | 6 (1) | 9 (0) | 4 (0) | 0 (0) |
|  |  | M | 8 (0) | 6 (0) | 2 (0) | 0 (0) |
|  | Malindi | F | 1 (0) | 35 (3) | 0 (0) | 0 (0) |
|  |  | M | 2 (0) | 17 (0) | 0 (0) | 0 (0) |
|  | Mombasa | F | 2 (0) | 23 (0) | 1 (0) | 0 (0) |
|  |  | M | 3 (0) | 10 (0) | 1 (0) | 0 (0) |
| Sub total |  |  | 22 (1) | 100 (3) | 8 (0) | 0 (0) |
| Grand total |  |  | 75 (13) | 116 (8) | 45 (4) | 23 (5) |

*Ae. aegypti* mosquitoes had 129 (60 females and 69 males) pools screened, 20.16% (n= 26) of the pools turned to be positive for flavivirus. There was site to site variation in terms of flavivirus positivity in the mosquito pools in three sites *(^χ2^, (df=2 n=30)* = 14.2292, P=0.001). In Kilifi, 18 pools (5 for females & 13 for males) of *Aedes aegypti* mosquitoes tested positive for flavivirus. In Mombasa, only three pools were positive comprising of *Aedes aegypti* mosquitoes only (1 pool for female and 2 for males). In Malindi, five pools of *Aedes aegypti* mosquitoes tested positive (all female pools). There was no significant difference between male and female positivity (χ^2^, *(df=1 n=30)* = 0.2697, P=0.604) (Table 6).

*Culex quinquefasciatus* mosquitoes had only 4 pools which tested positive, 1 pool in Kilifi, 3 in Malindi and none from Mombasa (Table 6). All flavivirus positive samples were negative for dengue virus.

## Discussion

Diverse ecological habitats were reported outdoors, though limited in numbers and corroborates Ngugi and others (51) results on the distribution of the breeding habitats. Discarded tyres, drums, water tanks, buckets, small domestic containers, water trough and Jerican has been identified to be the key breeding areas of *Aedes aegypti* mosquitoes (8,17,76, 77). Water storage containers (including Jericans, water tanks and drums) produced most of the immatures recorded, underscoring the importance of such containers in these regions. All larvae collected from indoor containers resulted in the emergence of only *Aedes aegypti* mosquitoes and this was consistent with other studies (10,76). Low indoor productivity in our study sites can also be attributed to human activities related to the use of domestic water storage devices. Most indoor containers are commonly used for hygiene, cooking and drinking and are subjected to frequent emptying and cleaning which can effectively interrupt mosquito development. They are therefore less likely to harbor mosquito immatures(76,78,79). In addition to this, most of the indoor containers for water storage were often covered; this could have possibly contributed to many of them being unproductive. Water-holding containers that are in frequent use within the domestic environment were observed to be less likely to harbor mosquito immatures (85) and this can make water storage possible without necessarily creating breeding sites for mosquitoes. Majority of the residents engage in small businesses which leads to indiscriminate disposal of waste (plastics, husks, polythene bags), and small scale farming. Local garages for tri/bi/motor-cycle and vehicle has led to the poor disposal of unused tyres. The flat terrain has poor drainage system leading to water logging during rainy season. Poor domestic and commercial waste disposable mechanisms and uncovered septic tanks and sewerage systems are common in the urban areas. Fewer roads are tarmacked, characterized by both covered and uncovered drainage systems along the streets. All these conditions, due to intensive and unplanned urbanization which results in largely modified topography and vegetation, provide suitable and numerous habitats for arboviral mosquito. Tires, which are important breeding sites for *Ae. aegypti* (3) produced only a small percentage of the larvae collected outdoors. This could be attributed to the period of collection as we were interested to establish the dry ecology of mosquitoes. Water tanks and small containers were mostly found in construction sites and commercial flower gardens as water holding containers. This shows that water tanks are suitable breeding habitats for mosquitoes in both indoors and outdoors. During the long dry season, in particular, drums and water tanks become important producers of *Ae. aegypti* immatures, if improperly covered, as they are used to store water, and this was consistent with other studies (51).

Furthermore, this entomologic investigation was also based on larval infestation indices (i.e. House index (HI), Container index (CI) and Breteau index (BI)). The Pan American Health Organization (PAHO) and World Health Organization (WHO) have described threshold levels for dengue transmission as low HI<0.1%, medium HI 0.1%—5% and high HI>5%(83)(86). The water storage practices could have resulted in high larval indices. In all the sites, these larval indices exceeded the WHO documented thresholds for risk of dengue outbreak/transmission, suggesting that all the areas sampled are at risk of dengue and other arboviral transmissions as reported in similar studies conducted along the coastal region documented high indices(10,76). Mombasa was at a higher risk of arboviral infections due to the higher larval indices followed by Kilifi and the least was Malindi.

Soil samples collected from the different potential habitats further shows that *Aedes* species can remain dormant for a long period in the soil as reported earlier in other studies(55). This container-breeding mosquito is well adapted to urban environments due to its preference for ovipositing in both natural and artificial water-filled receptacles, in which the nature of seasonally fluctuating water content leads to exposure of the eggs to drying conditions. The soil samples collected in this study were mostly from tyres. This further supports the preference of *Aedes* mosquitoes to oviposit in tyres(10,76). Previous study revealed that the desiccation resistance of *Aedes aegypti* eggs can be approximately 1 year, with another recent study showing almost the same results(83,84). The current study further reveals the importance of egg desiccation period since before the sampling was done, the study area had not experienced rains for a period of more than 8 months. Further studies on egg survivorship in soils are recommended to provide an understanding of the extent of *Ae. aegypti* intraspecific egg-desiccation resistance. This may allow a more refined modeling and provide greater insight into the current global distributions of the species, vector competence, and the intraspecific heterotic potential of more desiccation-resistant forms in the context of climatic stress or change. The unanticipated emergence or re-emergence of arboviral disease in recent years highlights the limits of our understanding of the dynamics that govern transmission of arboviruses. Without sufficient monitoring and surveillance programs to understand better the ecology of arboviral vectors, we will remain unprepared to prevent future epidemics from both unknown and known arboviruses.

The present study showed that potential arbovirus mosquito vectors are abundant and well distributed throughout the coastal region, although in varying densities. Variation in arboviral mosquito density and species richness was observed in the three sites, and could be due to the observed differences in the diversity of aquatic habitats among the three sites. Kilifi had more productive habitats compared to Malindi and Mombasa, thereby supporting diverse mosquito species. Previous studies (10, 76,85,86) have reported a positive relationship between habitat type diversity and mosquito species richness. *Aedes aegypti* and *Culex quinquefasciatus* are the most abundant mosquito species in urban areas of coastal Kenya, indicating that, potential arboviral vectors could be contributing to the current outbreaks of dengue, rift valley fever and Chikungunya in the coast region. Abundant *Cx. quinquefasciatus* mosquitoes were sampled across all the study villages, though in different densities. The most likely reason for this could be much of human activities which create microhabitats suitable and potential breeding areas of this species, such as; partial closed or open septic tanks, drainages, abandoned/unused swimming pools and drainage chambers (32,87,88). Flavivirus genus of arboviruses was isolated from *Culex quinquefasciatus* mosquitoes further supporting that these mosquitoes are vectors of an array of arboviruses and is consistence with other studies(3,80). *Aedes aegypti,* which is the principal vector of dengue virus, rift valley fever virus, Chikungunya, and urban yellow fever virus, represented 25.7% of the total collection and was the second predominant species. This species was well distributed in all the sampling sites though in varying densities and this implies that *Ae. aegypti* mosquitoes are well established in Coastal, Kenya, and the risk of Dengue fever, Chikungunya and yellow fever transmission would be high in the absence of effective vector control(3). Viruses were isolated from male mosquitoes suggesting that there could be trans-ovarian /vertical transmission of viruses from parents to offspring through the eggs. A similar study conducted in the coastal Kenya during the outbreak of dengue virus in 2013, isolated dengue virus from a pool of male *Aedes aegypti* mosquitoes (3). Reports of trans-ovarian transmission of dengue virus in *Aedes aegypti* mosquitoes have been reported in this region, therefore a lot of entomological surveillance should be conducted in this region to evaluate the extent, distribution and epidemiological significance of these viruses in the local vector populations. Phleboviruses and alphaviruses were negative in the study samples. Similarly, all flavivirus positive were specifically screened for dengue virus, and none of the pools turned positive. This does not mean that viruses of these genera are not circulating in the region. None of these mosquitoes was found to be positive for these viruses.

Mosquito (vector) blood-feeding patterns are important components of vector-borne disease transmission. The current study provides important information in estimating the degree of human-vector contact essential in understanding the role of a given species in disease transmission cycle. This study showed a majority of the blood meals were from humans, to lesser extent to chicken and the rest were unknown. Other animals present in the study area were dogs, cats, wild birds, and rodents, although logistical and resource limitations restricted ELISA tests against them despite these animals being known to be important blood meal sources for mosquitoes (97). It was shown that none of the mosquitoes had fed on goats or bovine although they are few or absent in the urban settings. All mosquito species tested showed low preference to human blood meal. *Culex quinquefasciatus* showed preference for both human and chicken whereas *Aedes aegypti* preferentially feed on human beings (81,35). Being vectors of many arboviruses, these mosquito species could be playing a significant role in the transmission of arboviruses in the coastal region of Kenya.

In conclusion, domestic and peri domestic containers were identified to be the key breeding areas of arboviral vectors. Therefore, efforts should be put in place targeting the productive habitat types. This study provides more information on arbovirus vectors distribution throughout the coastal region of Kenya in regions with previous history of outbreaks and where transmissions have not been reported. This highlights the potential for emergence of arboviruses in the coastal populations. Therefore, there is a need to map countrywide distribution and abundance of the Culicine mosquitoes beyond what this study has accomplished and conduct vector competence and blood meal assays for a comprehensive assessment of arbovirus risk to public health in Kenya. This will help to institute focused vector control measures in the event of a predicted outbreak. Of great importance, though, is the need to enhance surveillance activities for important arboviruses in livestock and humans and expand the prospective entomologic studies across the country.

### Ethics approval and consent to participate

The study was granted ethical approval from Kenya Medical Research Institute (KEMRI), Scientific and Ethics Review Unit (SERU), (Protocol SSC 2675) and the Pwani University Ethic Review Committee (ERC/MSc. /041/2016) prior to commencing of the research work. Oral informed consent was obtained from household heads to allow the survey of all accessible water-holding containers and setting up of traps in their residences.

### Consent for publication

All the authors have reviewed and approved the publication of this paper. This paper has been published with the permission of the Director of the Kenya Medical Research Institute (KEMRI).

### Availability of data and material

The supporting data is under the custodianship of the KEMRI-Wellcome Trust Data Governance Committee and is accessible upon request addressed to that committee.

### Financial support

This work was supported through the DELTAS Africa Initiative [DEL-15-003]. The DELTAS Africa Initiative is an independent funding scheme of the African Academy of Sciences (AAS)’s Alliance for Accelerating Excellence in Science in Africa (AESA) and supported by the New Partnership for Africa’s Development Planning and Coordinating Agency (NEPAD Agency) with funding from the Wellcome Trust [107769/Z/10/Z] and the UK government. The views expressed in this publication are those of the author(s) and not necessarily those of AAS, NEPAD Agency, Wellcome Trust or the UK government’. This work also received partial sponsorship on training of Culicine identification form KEMRI Internal Research Grant INNOV/IRG/020/2

### Authors’ contributions

JK, SMM, JM, CM conceived this study and drafted the first version of this manuscript. JK, JM, SMM, BK and CM conducted the field surveys, data collection and analysis. JK, DO, DN, MR and GW participated in the molecular screening of arboviruses CM offered scientific advisory leadership and reviewed/critiqued this manuscript. All of the authors have read and approved the final manuscript.

## Acknowledgement

This work formed part of the requirements for the Master’s degree of Pwani University. We are grateful to the Scientific and technical teams at the Centre for Geographic Medicine Research Coast, Kilifi for their contribution in the design and implementation of this work. Many thanks to the technical and field staff team of Festus Yaa, David Shida, Gabriel Nzai, Robert Mwakesi and Martha Muturi who devoted their time. They assisted in the field collection of mosquito samples and rearing in the insectary. Mr. Danstone Beti and Mr. John Gachoya (KEMRI-Center for Virus Research, Nairobi) who assisted in taxonomic or morphological identification of the mosquitoes and training of KEMRI CGMRC team on Culicines Morpho-taxonomy. We acknowledge Mr. Christopher Nyundo of KEMRI/Wellcome Trust Research Program in Kilifi for assisting in developing study area map.

